# Construction of recombinant *Bacillus subtilis* spores surface-displaying PRRSV ORF1b, GP5, M, and N proteins in tandem and evaluation of their immunogenicity in mice

**DOI:** 10.1101/2025.07.01.662551

**Authors:** Yang Yang, Jianzhen Li, Pengfei Fang, Bin Chen, Yicen Tang, Xixi Dai, Lei Fei, Yunfei Xiao, Yanpeng Dong, Chunfeng Shi, Xueqin Ni, Bo Jing, Yan Zeng, Kang cheng Pan

## Abstract

Porcine reproductive and respiratory syndrome virus (PRRSV), a single-stranded RNA virus, is a highly contagious pathogen that causes severe reproductive and respiratory disorders in pigs, leading to significant economic losses in the swine industry worldwide. However, current commercial vaccines provide only limited protection against circulating PRRSV strains, highlighting the urgent need for novel vaccine strategies. *B. Bacillus subtilis*, a well-characterized probiotic, has emerged as a promising platform for mucosal vaccine delivery due to its safety profile and ability to induce robust immune responses. In this study, we screened B cell and T cell linear epitopes from PRRSV-ORF1b, GP5, M, and N proteins to construct a recombinant tandem antigen (ER), which was displayed on the surface of *B. subtilis* 168 spores. The resulting recombinant strain, designated *B. subtilis* BE, was evaluated for immunogenicity in mice via oral administration. The results showed that Oral immunization with *B. subtilis* BE significantly elevated antigen-specific secretory IgA levels in intestinal contents and IgG levels in serum, indicating potent mucosal and systemic humoral responses. Moreover, *B. subtilis* BE induced measurable neutralizing antibody titers and enhanced cellular immunity, as evidenced by increased frequencies of CD3+ and CD8+ T cells in mesenteric lymph nodes and upregulation of *IFN-α, TNF-α*, and *IFN-γ* expression in intestinal tissues. Collectively, these findings demonstrate that *B. subtilis* BE elicits both humoral and cellular immune responses and may serve as a novel oral vaccine candidate against PRRSV, offering a promising alternative for controlling the ongoing epidemic.

## Introduction

Porcine reproductive and respiratory syndrome virus (PRRSV) is one of the most economically important pathogens affecting the global swine industry, especially in China and the United States[1]. PRRSV infection causes reproductive and respiratory disorders at various stages of pig development, characterized by respiratory symptoms in piglets and reproductive failure, fetal death, and congenital infection in sows[2]. The disease is identifiable by the distinctive blue-purple discoloration of the ears in affected pigs, commonly known as “pig blue ear disease”[3]. PRRSV is a single-stranded, positive-sense, enveloped RNA virus, with a genome size of 15kb, encoding at least ten open reading frames (ORFs): ORF1a, ORF1b, ORF2a, ORF2b, ORF3, ORF4, ORF5, ORF6, and ORF7[4]. PRRSV belongs to the order *Nidovirales* and the family *Arteriviridae*, emerging almost simultaneously as two species (*Betaarterivirus suid* 1 and *Betaarterivirus suid* 2), and with almost 50%– 70% nucleotide homogeneity[5]. The high genetic variability of PRRSV poses a major challenge for disease prevention and control in the global swine industry.

Vaccination remains the primary strategy for controlling the spread of this virus. Various vaccine platforms are currently available, including live attenuated, inactivated, subunit, DNA, and vector vaccines; but only modified live virus (MLV) and killed virus (KV) vaccines are commercially licensed for PRRS control[6]. The primary advantage of KV vaccines is their safety profile. However, their ability to induce robust immune protection is limited[7]. They often requires adjuvant support to enhance efficacy[8,9]. Although MLV vaccines generally confer effective protection against homologous viruses and partial immunity to heterologous strains[10,11]. Nonetheless, safety concerns remain due to risks of reversion to virulence and recombination between vaccine and virulent field strains[12–14]. Importantly, the continuous evolution of PRRSV poses an ongoing challenge[15]. There is an urgent need to develop novel vaccines that ensure safety and provide protection against both known and emerging strains.

GP5 and M proteins are the major structural proteins of PRRSV virus, and the GP5-M heterodimeric complex plays a critical role in virus budding and invasion of host cells[16,17]. Therefore, GP5-M protein is frequently targeted in the design for the development of genetically engineered vaccines. As research into PRRSV has advanced, novel protein combinations are being explored. Studies have demonstrated that the coexpression of GP5 with GP3, GP4, or N as fusion proteins is capable of eliciting enhanced immunogenicity compared to the expression of GP5 alone[18–20]. Moreover, the multi-epitope vaccines represent a distinctive design paradigm predicated on the incorporation of multiple overlapping antigenic epitopes[21]. The presentation of viral immunogenic proteins in complex formations is essential for the efficient induction of specific and protective B- and T-cell responses, which are crucial for eliciting robust immunity against PRRSV[6]. GP5, along with M, N, and certain non-structural proteins like ORF1b, harbors key virus-neutralizing B cell epitopes and potential T cell linear epitopes[22]. Furthermore, designing vaccines that target conserved epitopes is beneficial for developing a universal vaccine to overcome viral mutations.

The mucosal surfaces constitute the largest interface between the body and the external environment, encompassing the oral cavity, respiratory and gastrointestinal tracts, ocular and auditory cavities, and the genitourinary tract[23]. PRRSV primarily enters the host via respiratory and reproductive mucosa and causes disease predominantly at these mucosal sites[24]. Therefore, the development of a protective mucosal vaccine against PRRSV represents a promising strategy for controlling PRRS outbreaks. Mucosal vaccines are expected to induce protective cellular and humoral responses at both mucosal and systemic levels[25]. Robust activation of mucosal adaptive immunity including the production of secretory antibodies and the recruitment of tissue-resident T cells can prevent pathogen invasion and subsequent infection and transmission [26]. Moreover, as mucosal vaccines are administered non-invasively without injections, they offer ease of use and are well-suited for large-scale immunization programs.

*Bacillus subtilis* (*B. subtilis*) is a Gram-positive bacterium recognized for its remarkable adaptability and high productivity [27]. It is classified as “generally recognized as safe” (GRAS) by the US Food and Drug Administration, and it has been proposed as a probiotic in both human and animal uses[28,29]. It has strong secretory capacity, simple cell wall structure, well-characterized genetic background and non-toxicity[30]. These spores can survive indefinitely in harsh environments, including desiccation, nutrient deprivation, chemical stress, enzymatic degradation, extreme temperatures and pH levels [30–32]. One of the most notable applications of *B. subtilis* spores is in surface display technology. They can be easily engineered to display heterologous antigen or protein on their surface by utilizing natural surface proteins as anchors [33,34]. Furthermore, *B. subtilis* spores are capable of inducing strong mucosal immune responses and regulating the balance of Th1 and Th2 immune responses upon entering the host[32,35]. Consequently, these properties render spores a promising mucosal vaccine candidate. In 2001, Isticato [36] successfully displayed the 459 amino acid residues at the C-terminal of tetanus toxin protein on the spore surface of *B. subtilis* for the first time and induced a specific immune response. At present, the surface display technology of *B. subtilis* has been successfully applied to the immunization of pathogens of diverse types, like Singapore grouper iridovirus (SGIV)[37], *Streptococcus suis*[38], Carcinogenic Human Liver Fluke[39], and Porcine Rotavirus[40].

In this study, we designed a novel antigenic peptide incorporating both B-cell and T-cell linear epitopes, based on the conserved gene sequences of the ORF1b, GP5, M, and N proteins of the PRRSV NADC30-like strain. This peptide was successfully displayed on the surface of *Bacillus subtilis* 168 spores. We then assessed the ability of the recombinant spores to induce PRRSV-specific humoral and cellular immune responses in a mouse model. Our vaccine design strategy and experimental findings offer valuable insights for the development of next-generation PRRSV candidate vaccines and may also serve as a reference for designing vaccines against other pathogens.

## Results

### Analysis, characterization and evaluation of ER sequences

To enhance cross-protection against diverse PRRSV strains, we aligned the ORF1b, GP5, M and N domain sequences from different PRRSV strains, confirming that the selected domain is highly conserved. As it shown in Fig 1A, the ORF1b region of ER exhibits up to 92.7% sequence homology among different strains, with GP5, M and N’ homology shown in Fig S1-S3. The ER gene sequence spans 1,041 bp (Table S2), encoding a recombinant protein of 346 amino acids (Fig 1B). The construct includes single copies of ORF1b and N, and tandem repeats of GP5 and M, a design strategy aimed at enhancing immunogenicity[49]. The three-dimensional structure of the ER protein, predicted using AlphaFold3 (https://alphafoldserver.com/), exhibits a repetitive looped architecture (Fig 1C).

**Fig 1.**
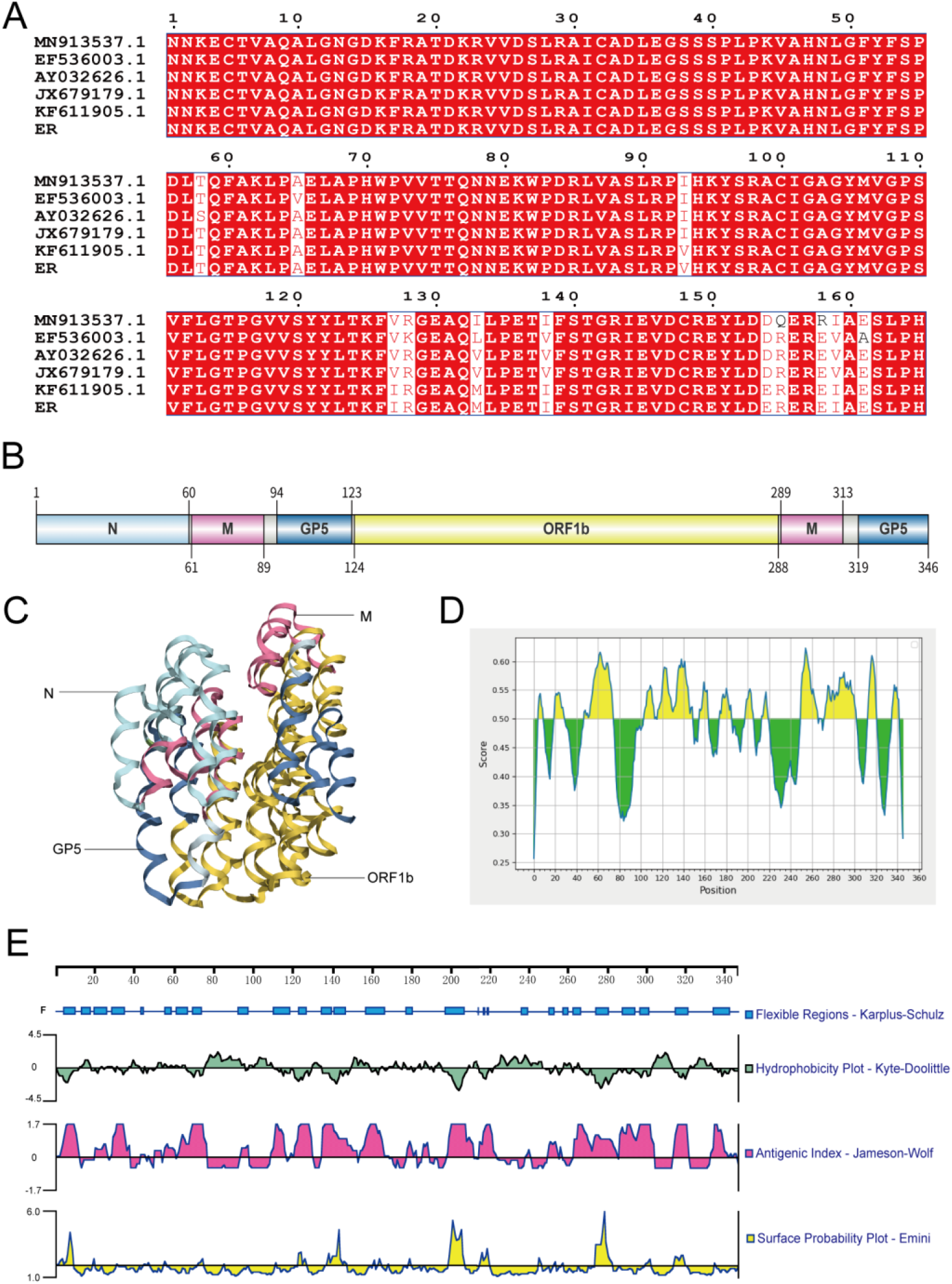
Analysis, characterization and evaluation of ER sequences. (A) Conservation analysis of the ORF1b region among different strains. (B) ER protein sequence. (C) The three-dimensional structure of the ER protein. (D) Prediction of B cell linear peptides by IEDB. (E) The results of ER protein predicted by Protean.

To evaluate its immunological potential, multiple computational tools were used. Fig 1D summarizes prediction scores at different locations on the ER’s structural proteins and shows the distribution of potential linear epitopes, within which several positions exhibits scores as high as 0.6. Following the T-cell epitope screening methods described by Zhou[50] and Sun[51], 21 peptide fragments with high immunoreactivity were identified. The specific B- and T-cell epitopes are listed in Tables S3 and S4, respectively.

Physicochemical properties of the ER protein, analyzed using DNAStar’s Protean software, are shown in Fig 1E. Most regions displayed a hydrophilicity index above zero, the protein demonstrates high hydrophilicity. The flexibility analysis shows that the protein has several flexible regions across a broad range. In terms of antigenic index, most regions have values greater than zero, suggesting a strong potential for antigenicity. The surface accessibility analysis indicates that these regions are exposed on the protein surface and are thus more likely to bind with antibodies and serve as antigenic epitopes.

### Prokaryotic expression of ER protein

As depicted in Fig 3A, the prokaryotic expression strain produced a protein band of approximately 45 kDa following induction, which aligns with the expected theoretical size. This confirmed that the ER protein could be efficiently expressed in a prokaryotic system.

### Construction of recombinant strain

The process of how to construct the recombinant spore is shown in Fig 2. The integrity of the recombinant plasmid pDG364-*CotB*-ER was confirmed by digestion with three distinct sets of restriction enzymes and Fig 3B illustrates that the recombinant plasmid was successfully constructed. In accordance with the principles of gene integration in *Bacillus subtilis* 168 mentioned above, the recombinant *Bacillus subtilis* lost its amylase activity, as demonstrated in Fig 3C. Furthermore, bacterial genomic DNA was validated through PCR using four specific primer pairs, yielding targeted bands at the anticipated locations, as displayed in Fig 3D. The band sizes of the PCR products using the genome of recombinant *B. subtilis* as a template were 4004 bp, 1041 bp, 2427 bp, and 2145 bp, respectively. In contrast, the PCR products of wild type *B. subtilis* had only one band of 569 bp when amyE-F/R were used as the primers, which proved that the size was consistent with the theoretical values. In summary, the recombinant *B. subtilis* was successfully constructed and we designated it as *B. subtilis* BE (Fig 3).

**Fig 2.**
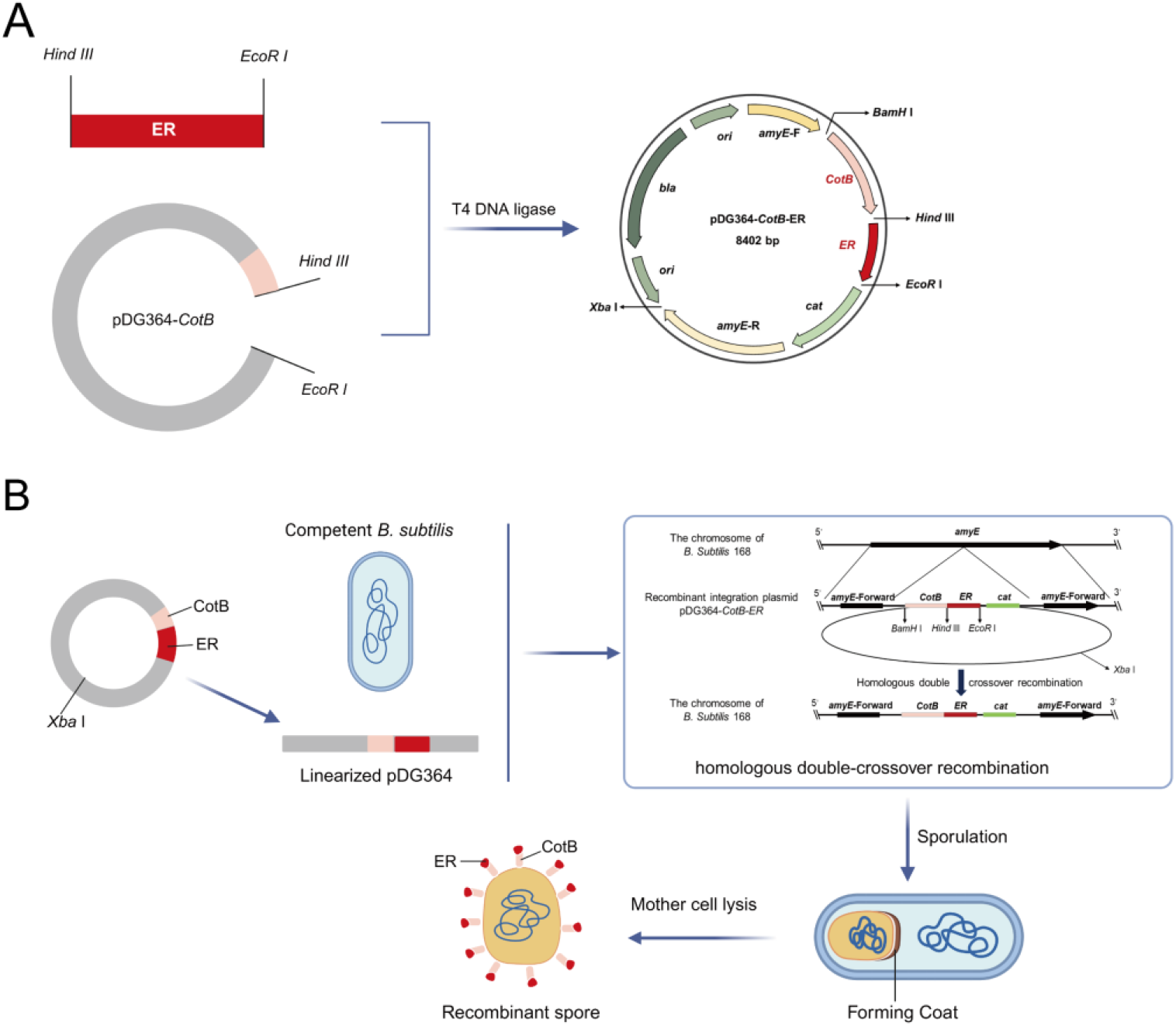
Process of spores surface display. (A) Construction of recombinant pDG364 plasmids. (B) Transformation of recombinant plasmids and display of outer coat of *CotB*-ER protein.

**Fig 3.**
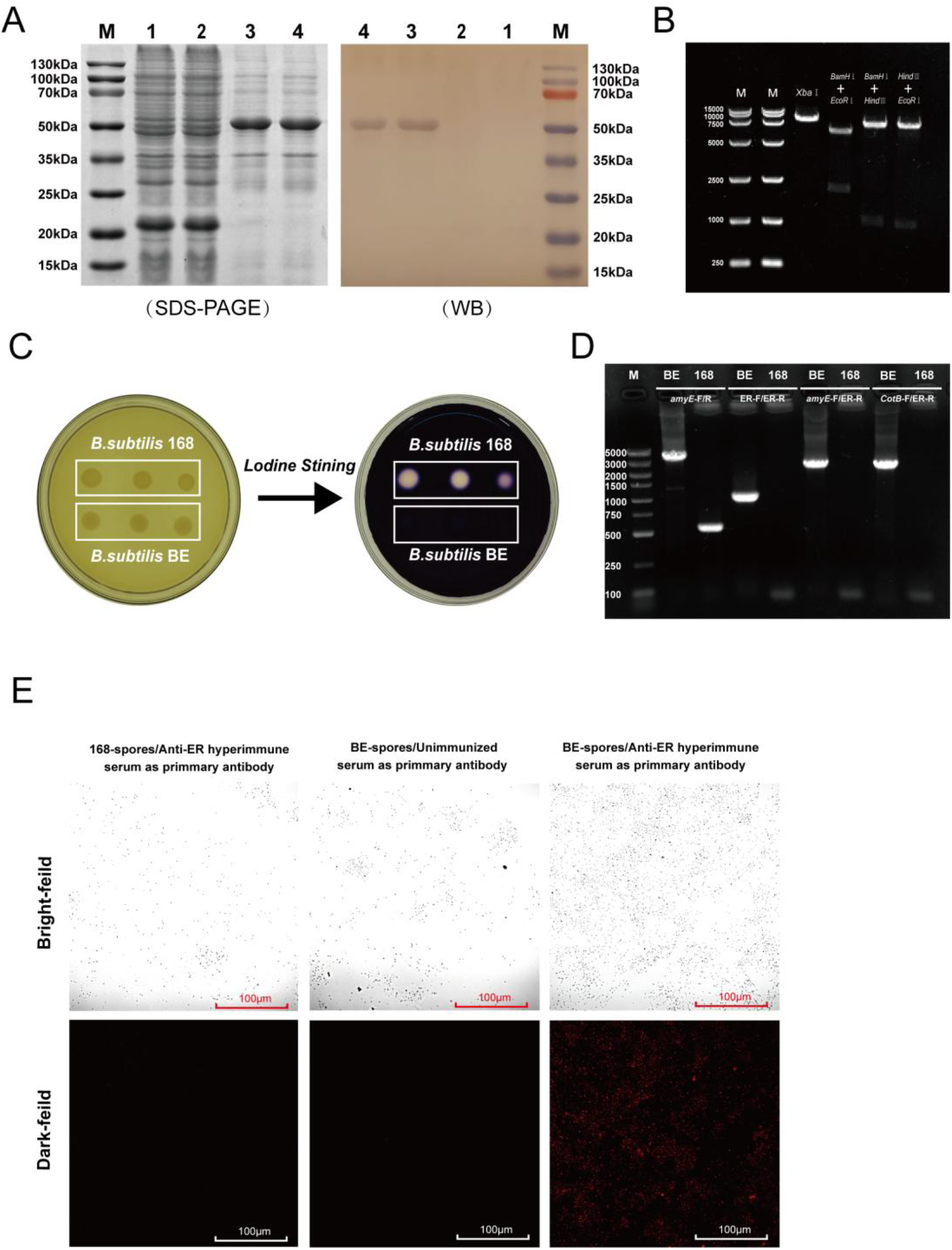
Verification of ER protein prokaryotic expression, recombinant pDG364 and recombinant *Bacillus subtilis* BE. (A) 12% SDS-PAGE and Western blot analysis of ER protein. Lane M: Protein marker (15–130 kDa); Lanes 1 and 2: Proteins expressed with the empty pET32a plasmid vector; Lanes 3 and 4: Expression products after induction with IPTG for 6 h. (B) The results of the recombinant plasmid pDG364-*CotB*-ER through double digestion. (C) The assessment of amylase activity in *B. subtilis* BE. (D) PCR verification of *B. subtilis* BE. (E) Indirect immunofluorescence assay results. The spores were observed under a 400× microscope. The scale bars represent 100 micrometers.

### Immunofluorescence detection

ER hyperimmune serum and Cy3-conjugated rabbit anti-mouse IgG were used respectively as primary antibody and secondary antibody for indirect immunofluorescence assay (Fig 3E). The results showed that only the spores of *B. subtilis* BE emitted fluorescent signal. In comparison, no specific fluorescence was detected when using serum from non-immunized mice as the primary antibody or when employing 168-spores as the carrier. These results confirm the successful presentation of the ER protein on the spore surface and its capacity to engage with specific antibodies, demonstrating *in vitro* reactogenicity.

### Detection of ER-specific antibodies by indirect ELISA

Fig 4A and Fig 4B presents the temporal dynamics of serum IgG and intestinal sIgA levels across different experimental groups over the feeding period. The ER group displayed significantly higher antibody levels compared to other groups as early as day 14 (*p*<0.0001), with mean P/N values of 2.03 for serum IgG and 3.11 for intestinal mucosal sIgA. This trend continued with a progressive increase in antibody response, reaching its peak at day 42. In contrast, intraperitoneal injection of commercial inactivated vaccines could not induce effective intestinal mucosal immunity in mice. Although the VR group exhibited higher levels of IgA antibodies on days 28 and 42 compared to the CK and WB groups (*p*<0.01), and showed a significantly elevated sIgA antibody concentration relative to the CK group at day 42 (*p*<0.01), these increases still remained negligible when compared to the ER group, suggesting that *B. subtilis* BE may provide long-lasting protection.

**Fig 4.**
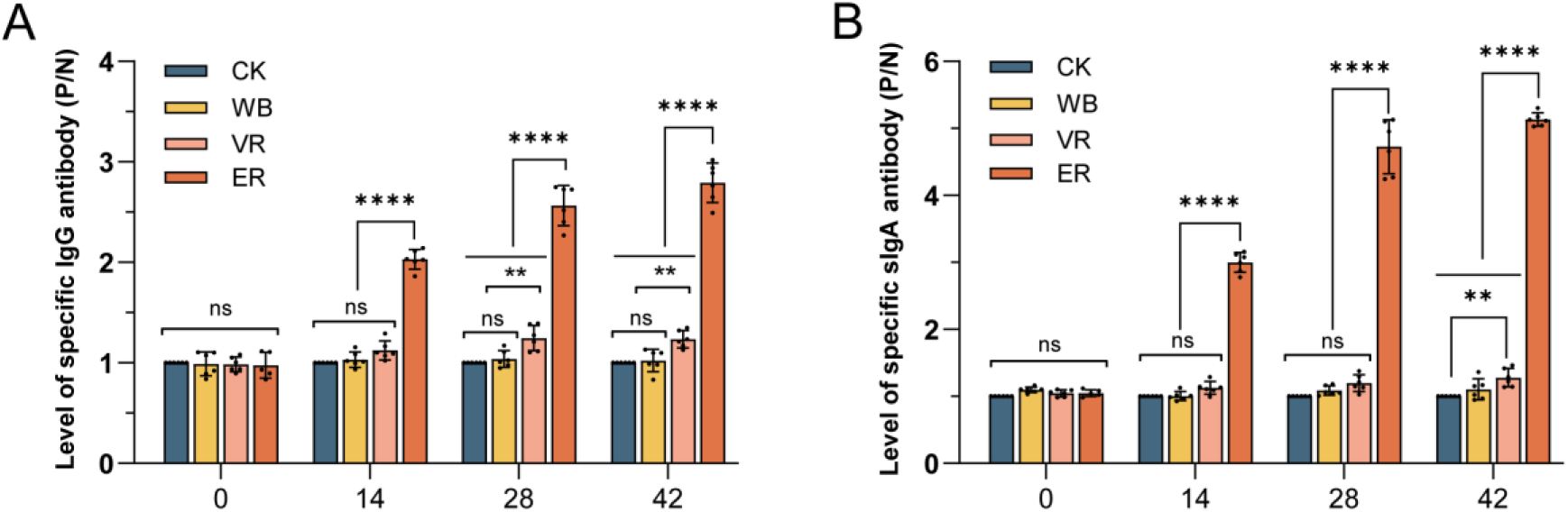
Detection of ER-specific antibodies by indirect ELISA. (A) Level of specific serum IgG antibody. (B) Level of specific intestinal sIgA antibody. Date are presented as mean ± SD with (*n* = 6). Statistical significance was calculated by two-way ANOVA with Tukey’s multiple comparisons test or Student’s t-test. Asterisks indicate significant differences (*, P < 0.05; **, P < 0.01; ***, P < 0.001; ****, P < 0.0001; ns, not significant).

### Virus microneutralization test

As shown in Fig 5B, the CK, WB and VR group did not produce any detectable VN antibodies specific for CH-1a. Only the ER group’s NA titers reached 1:4 (*p* < 0.0001). Although the NA titer is lower compared to that of the positive pig serum[52,53], it still demonstrates that BE-spores are capable of eliciting cross-neutralizing antibodies in mouse serum.

**Fig 5.**
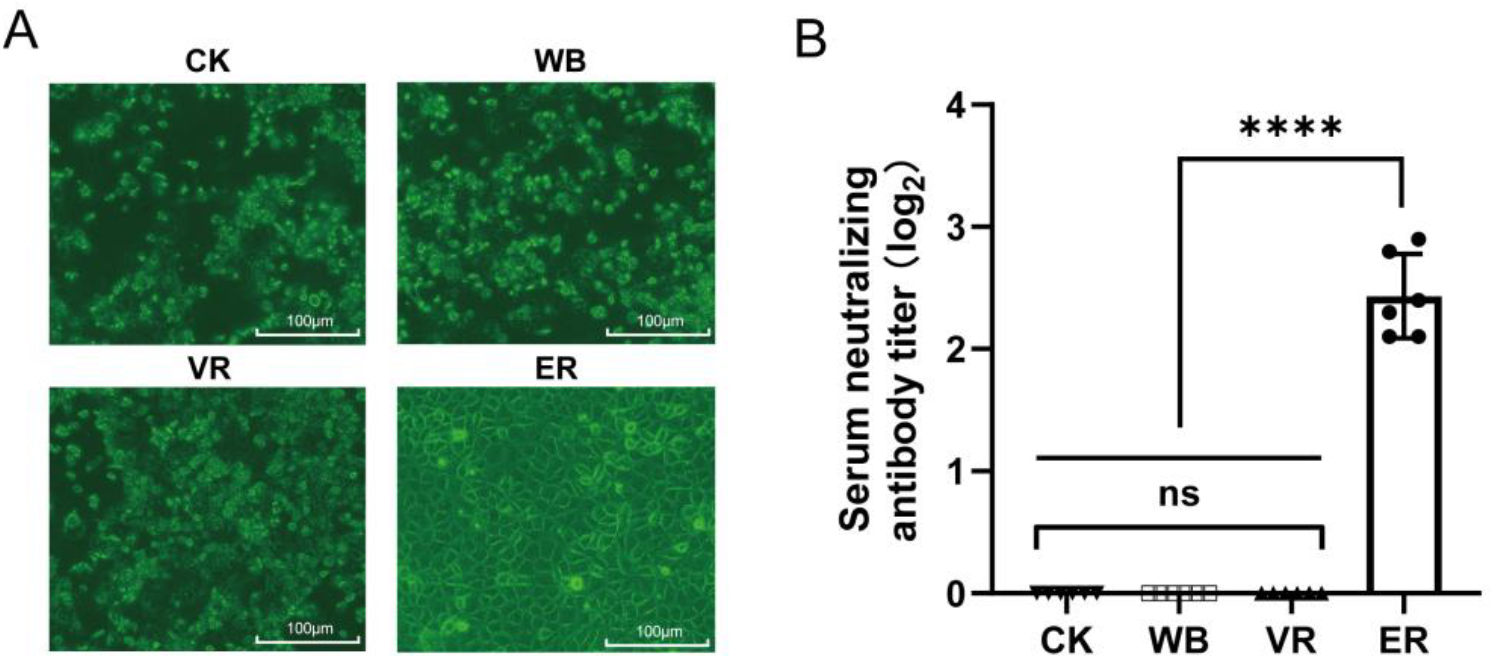
Results of virus microneutralization test. (A) Cytopathic effects (CPE) in Marc-145 cells. Serum from each group was co-cultured with the virus, subsequently infecting PRRSV strain R98, and after 72 hours the CPE was observed under a 400× microscope. The scale bars represent 100 micrometers. (B) The neutralizing antibody titers in mouse serum. Data are presented as mean ± SD (n = 6). Statistical significance was calculated by one-way ANOVA with Tukey’s multiple comparisons test or Student’s t-test. Asterisks indicate significant differences (*, P < 0.05; **, P < 0.01; ***, P < 0.001; ****, P < 0.0001; ns, not significant).

### Cytokine responses in immunized mice

Fig 6 shows that the mixed-feeding of BE-spores significantly enhanced the levels of *INF-α, TNF-α* and *IFN-γ* in mouse intestinal tissues (*p*<0.05). Meanwhile, it reduced the levels of Inflammatory cytokine *IL-6* (*p*<0.05). However, *IL-10* levels remained unaffected by the presence of BE-spores (*p*>0.05). As for the cytokine levels in serum, no significant differences were observed as is shown in Fig S5 (*p*>0.05), which demonstrates that the mucosal immunoadjuvant function by BE-spores is far more effective than the direct effect of spores to serum.

**Fig 6.**
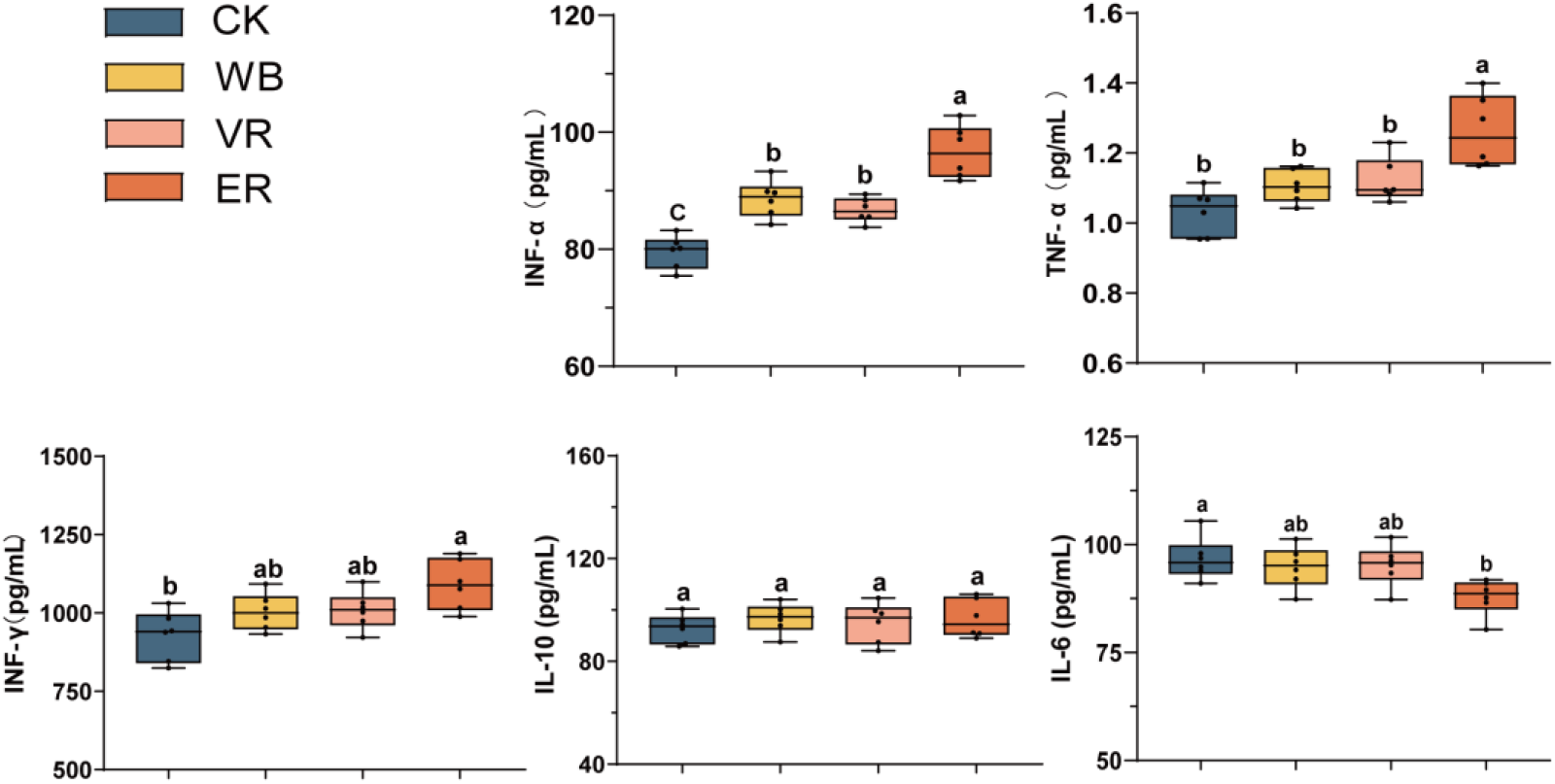
Levels of cytokines in mice ileal tissue measured by ELISA kit. Data are presented as mean ± SD (n = 6). Statistical significance was calculated by one-way ANOVA with Tukey’s multiple comparisons test or Student’s t-test. Bars with different letters are significantly different (*p* < 0.05). Bars sharing the same letter are not significantly different.

**Fig 6.**
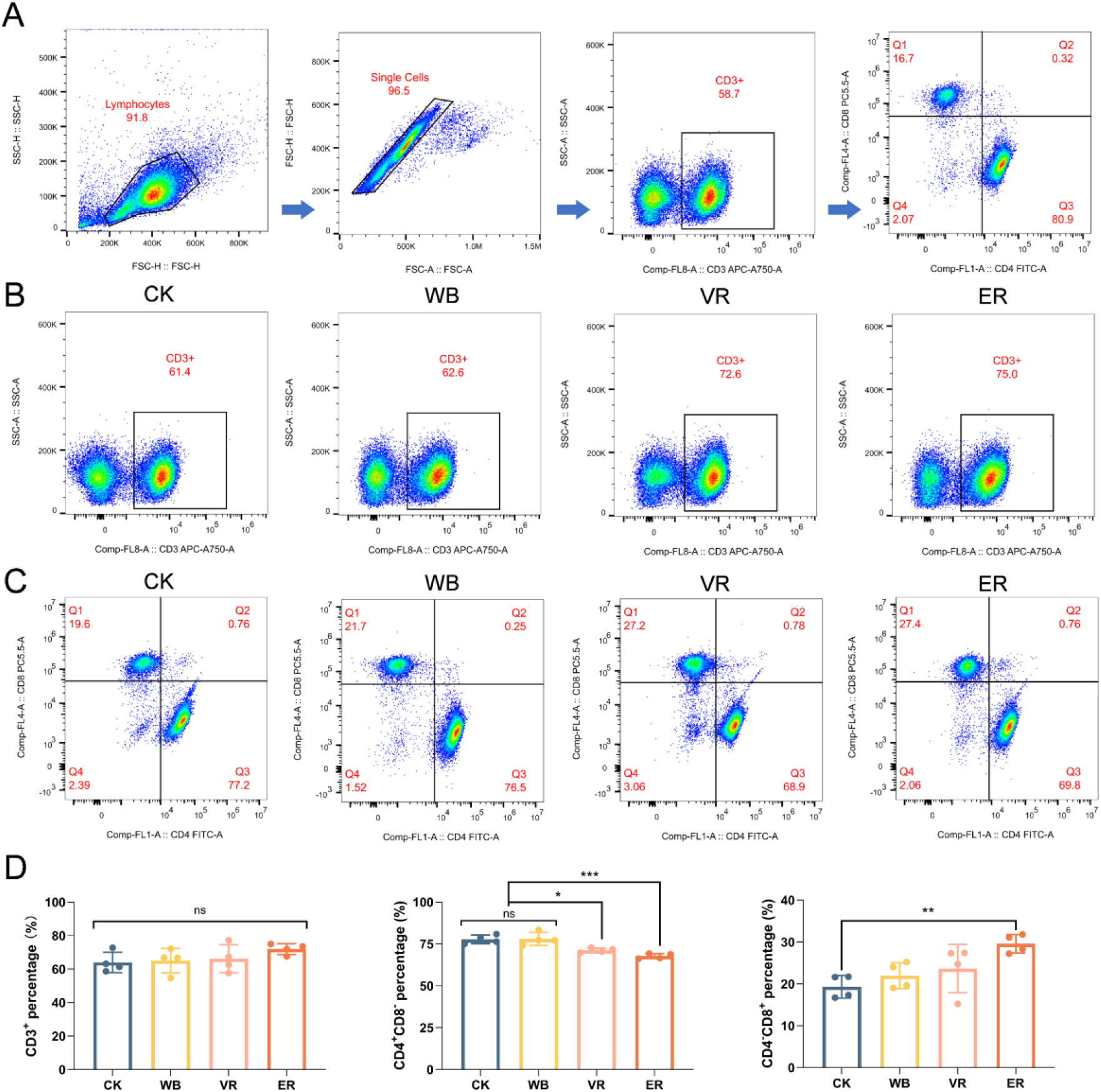
Detection of CD3^+^, CD4^+^ and CD8^+^ T lymphocytes in mesenteric lymph nodes (A) The gating strategy for flow cytometry applied to identify CD3^+^, CD4^+^ and CD8^+^ cell subsets. (B) The percentage of gated CD3^+^ T cells in the four experimental groups (C) The percentage of gated CD3^+^, CD4^+^ and CD8^+^ T cells in the four experimental groups. The CD4?CD8+ subset is located in quadrant Q1. The CD4^+^CD8+ subset is located in quadrant Q2. The CD4^+^CD8? subset is located in quadrant Q3. The CD4?CD8? subset is located in quadrant Q4. (D) The percentage of CD3^+^, CD4^+^ and CD8^+^ T subsets. Data are presented as mean ± SD (n = 4). Statistical significance was calculated by one-way ANOVA with Tukey’s multiple comparisons test or Student’s t-test. Asterisks indicate significant differences (*, P < 0.05; **, P < 0.01; ***, P < 0.001; ****, P < 0.0001; ns, not significant).

### Detection of CD3^+^, CD4^+^, CD8^+^ T lymphocytes in mesenteric lymph nodes

As illustrated in the gating strategy (Fig 6A), FSC-H was set as the X-axis and SSC-H as the Y-axis to initially gate the main lymphocyte population. Subsequently, FSC-A (X-axis) and FSC-H (Y-axis) were used to exclude cell doublets. Based on the use of Anti-Mouse CD3 antibody, 5μL of Anti-Mouse CD4 antibody, and 5μL of Anti-Mouse CD8a antibody, CD3 APC-A750-A was selected as the X-axis and SSC-A as the Y-axis to identify the CD3+ cell population. Then, CD4 FITC-A and CD8 PE-Texas Red (PT5.5-A) were used as the X-axis and Y-axis, respectively, to determine the proportions of CD4^+^ and CD8^+^ cells. Figs 6B and 6C display the representative gating of CD3^+^ T cell subsets and the distribution of CD4^+^/CD8^+^ subsets across groups. Quantitative comparisons of T cell subset proportions are shown in Fig 6D. No statistically significant differences in CD3^+^ T cell proportions were observed among the experimental groups (*p* > 0.05), although the ER group exhibited an upward trend. For CD4^+^ T cells, the VR group showed a significant reduction compared to both the CK and WB groups (*p* < 0.05), with a more pronounced decrease observed in the ER group (*p* < 0.001). In contrast, the proportion of CD8^+^ T cells was significantly increased in the ER group compared to the CK group (*p* < 0.01). Detailed numerical data for all T cell subsets are provided in Table S5.

## Discussion

PRRSV is a highly mutagenic RNA virus that continues to impose significant economic burdens on the global swine industry. Effective prevention and control are increasingly challenging due to the virus’s rapid evolution.[4]. Vaccination remains the primary strategy for controlling PRRSV. However, the current commercial vaccines suffer from critical limitations, including high dosage requirements, potential reversion to virulence, limited protection against homologous strains, and weak or no cross-protection against heterologous strains[10,11,54]. Given these limitations, novel alternative approaches are urgently required. *Bacillus*, a commonly used probiotic, is known for its ability to tolerate adverse conditions, including extreme temperatures and varying pH levels. It is widely utilized in livestock and aquaculture for its environmental resilience and gut-colonization capabilities. This capability enables *Bacillus* to exert significant probiotic effects and promote porcine growth[55,56]. Beyond that, its spore form can serve as highly effective delivery vehicles for the expressing foreign proteins. When administered orally, recombinant spores can survive passage through the digestive tract and interact with the mucosal immune system, making them attractive candidates for mucosal vaccine development.[57,58]. In this study, a recombinant *B. subtilis* strain was engineered to display a multiepitope fusion protein (ER) composed of PRRSV ORF1b, GP5, M, and N antigens on the spore surface. Oral immunization with these recombinant spores elicited strong mucosal and systemic immune responses in mice, including elevated levels of serum IgG, intestinal secretory IgA, and virus-neutralizing antibodies. Additionally, the spores stimulated increased intestinal expression of *IFN-α, TNF-α*, and *IFN-γ*, suggesting a robust Th1-type immune response. The expansion of CD8^+^ T cell populations in mesenteric lymph nodes further supports the induction of antigen-specific cellular immunity.

Secretory IgA (sIgA) plays a crucial role in the mucosal immune response, acting as the first line of defense against viral infections by preventing initial viral attachment and curbing systemic infection[59]. Our results demonstrated that oral administration of recombinant spores significantly enhances intestinal mucosal sIgA secretion in mice (Fig. 4B). The findings indicated that recombinant spores are capable of eliciting a robust mucosal immune response, which is consistent with the observations reported in previous studies on spore-based antigen delivery by Chen[60], Liang[61], Liu[47]. Moreover, the ER peptide retained its antigenicity when displayed on the spore surface, as evidenced by the high production of specific serum IgG (Fig 4A) and modest neutralizing activity (Fig 5). It is worth noting that the sera from mice immunized with the inactivated PRRSV vaccine exhibited no detectable virus-neutralizing titer, likely due to the low homology between the vaccine strain (CH-1a) and the challenge strain (R98). According to the findings of Castro[62] and Plaza-Soriano[63], the high heterogeneity of PRRSV might create differences in sensitivity to neutralization between isolates classified in heterogeneous categories. However, the ER antigen peptide was able to confer a certain degree of cross-protection, which is likely attributable to its derivation from a conserved region within the PRRSV genome. This finding underscores the importance of targeting conserved viral regions to enhance cross-protective efficacy, as demonstrated by the ER peptide.

The nature of the T-cell response is classified into either Th1 (*TNF-α* and *IFN-γ*) or Th2 (*IL-6, IL-10*) types of immune reactions, reflecting cellular and humoral immune responses, respectively[64]. *IL-10* inhibits the innate and adaptive immune responses from leukocytes and limits the potential tissue damage caused by inflammation[65]. Wang[66] reported a significant increase in *IL-10* expression following PRRSV vaccination. However, studies by Tsai[67] and Zhang[68] suggested that excessive or inappropriately timed *IL-10* production may facilitate PRRSV immune evasion during pathogenesis and potentially contribute to antibody-dependent enhancement (ADE). In our study, *IL-10* secretion levels did not differ significantly among the all groups, indicating that oral administration of recombinant spores is unlikely to promote ADE during PRRSV infection. *IL-6* is originally identified as a B-cell stimulatory factor and has important functions in regulating immune response, hemopoiesis, and inflammation. Xu[69] and Lothong[70] have suggested a potential correlation between *IL-6* expression levels and PRRSV virulence. Excessive expression of *IL-6* was relevant to the severe lung injury and damages of lymphoid organs. But in our experimental results, mice orally administered recombinant spores exhibited a significantly reduced level of *IL-6* in the intestinal mucosa (*p*<0.05). By contrast, elevated levels of *IFN-α, TNF-α*, and *IFN-γ* in the ER group may enhance antiviral immunity and promote viral clearance. As a result, based on the overall cytokine measurement results, oral administration of recombinant spores primarily promoted a Th1-type immune response, as evidenced by elevated intestinal secretion of *TNF-α* and *IFN-γ* observed in our study. This also aligns with the findings of Tobita, who reported that *Bacillus subtilis* BN could stimulate the secretion of Th1-type cytokines through interaction with Toll-like receptor 2 (TLR2)[71].

To assess the ability of *B. subtilis* BE to induce cellular immunity, flow cytometry was conducted to analyze the populations of CD3^+^, CD4^+^, and CD8^+^ cells in mouse mesenteric lymph nodes (MLNs). We selected mesenteric lymph nodes (MLNs) as the sampling site because MLNs are key components of the inductive sites of the gut-associated lymphoid tissue (GALT), which are essential for immune homeostasis.[72]. CD3^+^ T cells include both CD4^+^ helper T cells and CD8^+^ cytotoxic T cells. CD4^+^ cells regulate immune responses through cytokine secretion, while CD8^+^ cells directly kill infected or abnormal cells[73,74]. Fig 7D showed that there was no significant difference in the proportion of CD3^+^ cells in the MLNs among the groups (*p*>0.05). However, the ER group, which was administered recombinant spores, showed a trend toward an increased proportion(*p*>0.05). Besides, *B. subtilis* BE induced preferential CD8^+^ T cell proliferation and reduced CD4^+^ T cell proliferation. This could be attributed to the regulatory function of CD8^+^ T cells, which can exert immunomodulatory effects by eliminating CD4^+^ T cells[75]. In Mou’s research, the engineered mRNA vaccine encoding the full-length GP5 and M proteins (GP5-M) elicited both the proliferation of PRRSV-specific T cells and a marked expansion of CD8^+^ T cells as well[76]. Sun et al also demonstrated that the nanoparticle vaccine encoding GP3, GP4 and GP5 proteins triggered *IFN-γ* production and promoted the differentiation of CD8^+^ T cells[51]. These researches highlight the role of PRRSV-specific cytotoxic T lymphocytes (CTLs) in viral clearance, indicating that the antigen-specific cellular immunity extends beyond humoral immunity and contributes to protective immunity against PRRSV.

Despite promising findings, this study has several limitations. Due to technical challenges in spore protein extraction, western blot analysis of ER peptide was conducted only on prokaryotically expressed protein, rather than spore-derived proteins. Furthermore, the immunogenicity of individual epitopes within the ER fusion protein was not separately assessed. Future studies in pigs are essential to validate the vaccine’s protective efficacy in a natural host. Additionally, the cross-protective ability of *B. subtilis* BE against more different PRRSV variants has not been assessed, which is particularly important in the context of PRRSV’s high variability.

## Conclusion

In summary, we developed a novel multiepitope recombinant *Bacillus subtilis* strain (BE) that displays PRRSV ORF1b, GP5, M, and N antigens in tandem on the spore surface. Oral administration of *B. subtilis* BE in mice elicited strong antigen-specific mucosal and systemic humoral responses, as evidenced by elevated levels of secretory IgA and serum IgG. The recombinant spores also modulated cytokine expression in the intestinal mucosa and enhanced cellular immune responses, including increased CD3^+^ and CD8^+^ T cell populations. These findings collectively demonstrate the potential of *B. subtilis* BE as a promising oral vaccine candidate for the prevention of PRRSV infection.

## Materials and Method

### Strains, cell lines, vaccines, plasmids and primer sequences

The strains, cell lines, vaccines, plasmids and primer sequences used in this study are listed in Supplementary Table S1.

### Design and prokaryotic expression of ER peptides in *E coli* BL21

The strategy for selection of the PRRSV gene region was designed based on the type NADC30-Like isolate (Gene bank: KF611905.1). We employed DNAMAN 9.0 to identify conserved amino acid sequences within PRRSV ORF1b, ORF5, ORF6, and ORF7 regions. B and T cell epitopes were then predicted within these sequences using NetMHCpan 4.1 (https://services.healthtech.dtu.dk/services/NetMHCpan-4.1/) and IEDB (http://tools.iedb.org/mhci/). After evaluation, suitable fragments were concatenated to form a final composite fragment, designated as the ER gene (listed in Supplementary Table S2). The ER gene fragment was synthesized by Qingke Biotechnology and cloned into the pET32a vector (containing a His tag) after restriction endonuclease digestion with *Kpn I* and *XhoI*. The prokaryotic expression vector pET32a-ER was transformed into *E. coli* BL21 cells, and a single colony was inoculated into Luria–Bertani (LB) broth containing 50 mg/L ampicillin (Sigma, USA) and incubated at 37°C. Overnight cultures were transferred into 100 ml of fresh LB medium for large-scale protein production at 37°C. Expression of recombinant HR2P-His was induced with isopropyl-b-D-thiogalactoside (IPTG) (Sigma, USA) at a final concentration of 0.4 mM when Ab_s600 nm_ reached 0.5. After induction for 6 h at 37°C, the obtained precipitate after ultrasonic disruption was dissolved in 8 M urea and subjected to 12% SDS-PAGE, followed by Coomassie brilliant blue G-250 staining. After completion of the SDS-PAGE, the samples were transferred to a nitrocellulose membrane (Servicebio, Wuhan). Following overnight blocking with 2% bovine serum albumin (BSA) in TBST, the samples were incubated for one hour at room temperature with mouse anti-his tag antibody (1:1000 diluted in TBST, Cat#M30975-1, BOSTER, Wuhan). They were then washed with TBST and incubated for one hour at 25°C with horseradish peroxidase (HRP)-conjugated goat anti-mouse IgG (1:2000 diluted in TBST, Cat#BA1050, BOSTER, Wuhan) as the secondary antibody. Finally, the recombination ER protein carrying the His tag appeared as brown bands when stained with DAB Substrate Kit (BOSTER, Wuhan).

### Construction of recombinant integration plasmid

PCR amplification of the pET32a-ER construct was performed using ER-F2/ER-R2 primers, followed by the original *Kpn I* and *Xho I* restriction sites being replaced with *Hind III* and *EcoR I*. The pDG364-*CotB* plasmid was double-digested with *Hind III* and *EcoR I* restriction enzymes, and the purified PCR product (TIANGEN, Beijing) was ligated into the vector using T4 DNA ligase (Beyotime, Shanghai), in accordance with our previous study[40]. The correctly constructed plasmid, confirmed by double digestion, was designated as pDG364-*CotB*-ER.

### Construction and verification of recombinant *B subtilis* 168

Competent cells of *B. subtilis* 168 were prepared according to the method of Julkowska[41]. To enhance homologous recombination efficiency, the pDG364-*CotB*-ER plasmid was linearized with the restriction enzyme *Xba I*[42]. Added an appropriate amount of linearized pDG364-*CotB*-ER into 500 μL of competent cells, mixed gently and shaked slowly at 37°C (80 r/min) for 1 h. The integrated plasmid pDG364-*CotB*-ER employs the upstream and downstream regions of the *B. subtilis* amylase gene as homologous arms. Upon transformation of plasmid pDG364-*CotB*-ER into *B. subtilis* 168, homologous double-crossover recombination occurred between the plasmid’s homology arms and the corresponding chromosomal regions under antibiotic selection. Positive clones were initially screened on LB agar containing 5 mg/L chloramphenicol and on 1% starch nutrient agar. Starch plates were stained with iodine solution to visualize clear hydrolysis zones. Recombinant strains lacking amylase activity were selected, and their genomic DNA was extracted using the Bacterial Genomic DNA Isolation Kit (Sagon, Shanghai). Wild-type *B. subtilis* 168 served as the control. PCR and sequencing were conducted using specific primer pairs (Supplementary Table S1). The strain with correct genomic integration of the ER gene was designated *B. subtilis* BE.

### Preparation of spores and indirect immunofluorescence

Used Difco Sporulation Medium (DSM) to sporulate the *B. subtilis* BE and *B. subtilis* 168 according to the nutrient depletion method[43]. The purified spores were fixed and observed by immunofluorescence microscopy to identify whether the ER protein was successfully displayed on the spore surface. The primary antibody was mouse-derived anti-ER hyperimmune serum as a positive control (Diluted to 1:100 in PBST), and the secondary antibody was fluorescein Cy3-conjugated goat anti-mouse IgG (Diluted to 1:100 in PBST, Biomed, Beijing). Also, serum from unimmunized mice was used as a negative control, then indirect immunofluorescence of *B. subtilis* BE spores were performed with Cy3-conjugated goat anti-mouse IgG. Finally, samples were detected under a onfocal laser scanning microscope(CLSM)(OLYMPUS, FV4000, Japan).

### Oral immunization of mice and sample collection

The Institutional Animal Care and Use Committee at the Sichuan Agricultural University approved all the procedures used in this study. Approval number was SYXK Chuan 20250030. The study methodologies were complied with the Guide for the Care and Use of Laboratory Animals and utmost care was taken to reduce animal discomfort. Ninety-six female BALB/c mice, aged 3 weeks, along with basal diets, were procured from Chengdu Dossy Experimental Animal Co., Ltd. Following a 7-day acclimatization period, the mice were randomly divided and assigned to four groups (24 mice per group). *B. subtilis 168* and *B. subtilis* BE were fermented and sporulated using DSM, and the number of spores was adjusted to immunize mice according to the immunization program in Table 1. The nutritional composition of the mouse diet met the standard growth requirements, and the number of spores added referred to Dai, Chen, Li and Liu[44–47]. Each of the mice were fed 3-5 g per day. Euthanized 6 mice of each group at the set time for sampling. Serum, intestinal contents and tissues were collected on days 0, 14, 28, and 42. All samples were then stored at -80°C. In addition, fresh intestinal lymph nodes were collected from mice on day 42 and preserved in sterile PBS on ice for subsequent flow cytometry analysis.

**Table 1.** Program for vaccination experiment

| Group | Strains | Method of Administration | Dose | Time of Administration |
| --- | --- | --- | --- | --- |
| CK | — | — | — | — |
| WB | *B. subtilis* 168 spores | Fed the mixed diet | $2.0 \times 10^6$ CFU/g feed | The whole period |
| VR | Inactivated vaccine | multi-site subcutaneous injection | 100 μL /per mouse | Days 1, 14, and 28 |
| ER | *B. subtilis* BE spores | Fed the mixed diet | $2.0 \times 10^6$ CFU/g feed | The whole period |

### Serum IgG and intestinal content IgA detection

According to the method described by Zhang et al.[48], purified ER protein (0.5 μg/mL) in 0.05 M carbonate buffer (pH 9.6) was coated onto ELISA plates (Servicebio, Wuhan) at 100 μL per well. After incubation overnight at 4°C, the plates were blocked with 200 μL PBST containing 2% BSA and incubated at 37°C for 2 hours. Following three washes with PBST, 100 μL/well of serum and small intestine content samples, diluted to 1:500 with PBST, were added and incubated for 1 hour at 37°C. HRP-conjugated goat anti-mouse IgG (diluted 1:4000 in PBST) and HRP-conjugated goat anti-mouse IgA (diluted 1:30,000 in PBST, Abcam, Cat#ab97235, Cambridge) were used as secondary antibodies and incubated at 37°C for 30 minutes. The reaction was developed with 100 μL TMB substrate (Sangon, Shanghai) in the dark for 10 minutes. The reaction was stopped by adding 50 μL 2 M H_2_SO_4_, and absorbance was measured at 450 nm using a microplate reader (Varioskan Flash, Thermo Fisher Scientific, Waltham, MA, USA). Serum IgG and intestinal mucosal secretory IgA (sIgA) levels were expressed as positive-to-negative (P/N) ratios, with the CK group serving as the negative control and P/N≥2.1 as positive.

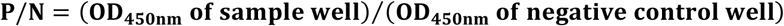

### Serum neutralization assay

The neutralizing antibody (NAb) titers in mouse serum were determined on Marc-145 cells. Briefly, the viruses were diluted to a concentration of 100 TCID50 per 50 μL in DMEM supplemented with 2% FBS. Serial dilutions of serum, collected at 42days post-inoculation and starting at a 1:4 dilution, were mixed with the diluted viruses and incubated at 37°C for 1h. Subsequently, 100 μL of each mixture was transferred to Marc-145 monolayers in 96-well plates and incubated for an additional 2 days at 37°C with 5%CO_2_. Cytopathic effects (CPE) were evaluated under light microscopy. A reduction of more than 90% in infected cells was considered indicative of neutralization at that dilution; otherwise, the sample was deemed negative for neutralization. Each serum dilution was tested in quadruplicate wells. NAb titers were calculated using the Reed-Muench method.

### Cytokine responses detection

A sample of 0.1 g of intestinal tissues, collected at 42 days post-inoculation, was weighed and mixed with 0.9 mL of PBS. The mixture was centrifuged at 3000 rpm for 10 minutes to isolate the supernatant. ELISA kit (Jiancheng Bioengineering Institute, Nanjing) was used to measure the levels of cytokines *INF-α, TNF-α, INF-γ, IL-10* and *IL-6* in supernatant and serum according to the manufacturer’s instructions.

### Flow Cytometry

Fresh mesenteric lymph nodes, collected at 42 days post-inoculation, were placed on a 70 μm cell strainer and gently dissociated using a pestle. The resulting cell suspension was collected into a 50 mL centrifuge tube. Cells were centrifuged at 300 ×g for 5 min at 4 °C, and the supernatant was discarded. The pellet was resuspended in 1 mL ACK lysis buffer and incubated at room temperature for 5 min to lyse red blood cells. Lysis was terminated by adding 5 mL RPMI-1640 medium supplemented with 2% FCS, followed by vortexing. The suspension was filtered again through a 70 μm strainer and centrifuged at 300 ×g for 5 min at 4 °C. The supernatant was removed, and the pellet was resuspended in staining buffer at a final concentration of 1×10^7^ cells/mL. For surface staining, 100 μL of single-cell suspension was added to each flow cytometry tube, followed by 5 μL each of anti-mouse CD3, CD4, and CD8a antibodies. Samples were vortexed and incubated protected from light for 15-30 min at room temperature or 30–60 min on ice. Subsequently, 2 mL of PBS was added to each tube, vortexed, and centrifuged at 300 ×g for 5 min at room temperature. The supernatant was discarded, and the pellet was resuspended in 500 μL Flow Cytometry Staining Buffer for acquisition. Data were acquired by flow cytometry and analyzed using FlowJo v10.8.1.

### Statistical Analysis

Data were displayed as means ± standard deviation (SD) values. One-way analysis of variance (ANOVA) with Dunnett’s post-hoc test was used for multiple comparisons to determine significant differences between groups by using GraphPad Prism software (version 10.0). A *p*-value of less than 0.05 was considered to be statistically significant. Indications of statistical significance are represented as * for *p*<0.05, ** for *p*<0.01, and *** for *p*<0.001, **** for *p* < 0.0001; ns for not significant.

## Supporting information

**S1 Table: The strains, cell lines, vaccines, plasmids and primer sequences used in this study. (docx) S2 Table: ER gene sequences. (docx)**

**S3 Table: Results of B cell linear epitope prediction in ER. (docx) S4 Table: Results of T cell linear epitope prediction in ER. (docx)**

**S5 Table: The proportion of T cell subsets analyzed by flow cytometry. (docx)**

**S1 Fig: Multiple sequence alignment of GP5 protein. (docx)**

**S2 Fig: Multiple sequence alignment of M protein. (docx)**

**S3 Fig: Multiple sequence alignment of N protein. (docx) S4 Fig: PCR verification of ER gene. (docx)**

**S5 Fig: Levels of cytokines in mice serum at 42 days measured by ELISA kit**. Values are presented as mean ± SD with (n = 6). Statistical significance was calculated by one-way ANOVA with Tukey’s multiple comparisons test or Student’s t-test. Bars with different letters are significantly different (p < 0.05). Bars sharing the same letter are not significantly different. (docx)

### Experimental ethics

The Institutional Animal Care and Use Committee at the Sichuan Agricultural University approved all the procedures used in this study. Approval number was SYXK Chuan 20250030. And the study methodologies were complied with the Guide for the Care and Use of Laboratory Animals.

## Author contributions

**Conceptualization:** Yang Yang, Jianzhen Li, Pengfei Fang, Kangcheng Pan.

**Methodology:** Yang Yang, Yicen Tang, Lei Fei, Chunfeng Shi.

**Formal analysis and investigation:** Yang Yang, Bin Chen, Yanpeng Dong.

**Visualization:** Yang Yang, Yunfei Xiao.

**Writing-original draft preparation:** Yang Yang, Jianzhen Li, Xixi Dai.

**Writing-review an editing:** Xueqin Ni, Yan Zeng.

**Supervision:** Bo Jing, Yan Zeng.

**Resources:** Yan Zeng, Kangcheng Pan.

**Funding acquisition:** Kangcheng Pan.

## Funding

This work was supported by grants from the Technology Innovation Research Team in the University of Sichuan Province (Award Number: KM406183.1), and The School-Enterprise Cooperation Project of Sichuan Agricultural University (Award Number: 2024511825000383).

## Competing interests

The authors have declared that no competing interests exist.

## Data availability statement

All data generated or analyzed during this study are included in this published article.

## Notes

### Competing Interest Statement

The authors have declared no competing interest.

